# Fear in dreams and in wakefulness: evidence for day/night affective homeostasis

**DOI:** 10.1101/534099

**Authors:** V Sterpenich, L Perogamvros, G Tononi, S Schwartz

## Abstract

Recent neuroscientific theories have proposed that emotions experienced in dreams contribute to the resolution of emotional distress and preparation for future affective reactions. We addressed one emerging prediction, namely that experiencing fear in dreams is associated with more adapted responses to threatening signals during wakefulness. Using a stepwise approach across two studies, we identified brain regions activated when experiencing fear in dreams and showed that frightening dreams modulated the response of these same regions to threatening stimuli during wakefulness. Specifically, in Study 1, we performed serial awakenings in 18 participants recorded throughout the night with high-density EEG and asked them whether they experienced any fear in their dreams. Insula and midcingulate cortex activity increased for dreams containing fear. In Study 2, we tested 89 participants and found that those reporting higher incidence of fear in their dreams showed reduced emotional arousal and fMRI response to fear-eliciting stimuli in the insula, amygdala and midcingulate cortex, while awake. Consistent with better emotion regulation processes, the same participants displayed increased medial prefrontal cortex activity. These findings support that emotions in dreams and wakefulness engage similar neural substrates, and substantiate a link between emotional processes occurring during sleep and emotional brain functions during wakefulness.

**SIGNIFICANT STATEMENT:** Highly debated while pivotal to current theoretical models of dreaming, the relationship between emotion processing during wakefulness and in dreams remains elusive. In a first study, we used high-density EEG recordings and observed that regions involved in fear processing (i.e. the insula and midcingulate cortex) were activated during fear-related dreams. This first finding demonstrates that emotions in dreams engage similar neural circuits as during wakefulness. In a second study, using fMRI, we show that higher incidence of fear in dreams was associated with reduced emotional arousal and brain responses indicative of better emotion regulation during wakefulness. Together, these results strongly support the idea that experiencing fear in a secure environment, as in dreams, relates to more adapted responses to threatening events in real life.

## INTRODUCTION

Converging evidence from human and animal research suggests functional links between sleep and emotional processes (Wagner U et al. 2006; Walker MP and E van der Helm 2009; Perogamvros L and S Schwartz 2012; Boyce R et al. 2016). Chronic sleep disruption can lead to increased aggressiveness (Kamphuis J et al. 2012) and negative mood states (Zohar D et al. 2005), while affective disorders such as depression and PTSD are frequently associated with sleep abnormalities (e.g., insomnia and nightmares). Experimental evidence indicates that acute sleep deprivation impairs the prefrontal control over limbic regions during wakefulness, hence exacerbating emotional responses to negative stimuli (Yoo SS et al. 2007). Neuroimaging and intracranial data further established that, during human sleep, emotional limbic networks are activated (e.g., Maquet P et al. 1996; Braun AR et al. 1997; Nofzinger EA et al. 1997; Schabus M et al. 2007; Corsi-Cabrera M et al. 2016). Together these findings indicate that sleep physiology may offer a permissive condition for affective information to be reprocessed and reorganized. Yet, it remains unsettled whether such emotion regulation processes also happen at the subjective, experiential level during sleep, and may be expressed in dreams. Several influential theoretical models formalized this idea. For example, the Threat Simulation Theory postulated that dreaming may fulfill a neurobiological function by allowing an offline simulation of threatening events and rehearsal of threat-avoidance skills, through the activation of a fear-related amygdalocortical network (Revonsuo A 2000; Valli K et al. 2005). Such mechanism would promote adapted behavioral responses in real life situations (Valli K and A Revonsuo 2009). Other models suggested that dreaming would facilitate the resolution of current emotional conflict (Cartwright R et al. 1998; Cartwright R et al. 2006), the reduction of next-day negative mood (Schredl M 2010) and extinction learning (Nielsen T and R Levin 2007). Although these two main theories differ, because one focuses on the resolution of current emotional distress (e.g. fear extinction; Nielsen T and R Levin 2007) and the other on the optimization of waking affective reactions (Revonsuo A 2000; Perogamvros L and S Schwartz 2012), both converge to suggest that experiencing fear in dreams leads to more adapted responses to threatening signals during wakefulness (Scarpelli S et al. 2019). The proposed mechanism is that memories from a person’s affective history are replayed in the virtual and safe environment of the dream so that they can be reorganized (Nielsen T and R Levin 2007; Perogamvros L and S Schwartz 2012). From a neuroscience perspective, one key premise of these theoretical models is that experiencing emotions in dreams implicates the same brain circuits as in wakefulness (Hobson JA and EF Pace-Schott 2002; Schwartz S 2003). Preliminary evidence from two anatomical investigations showed that impaired structural integrity of the left amygdala was associated with reduced emotional intensity in dreams (De Gennaro L et al. 2011; Blake Y et al. 2019).

Like during wakefulness, people experience a large variety of emotions in their dreams, with REM dreaming being usually more emotionally-loaded than NREM dreams (Smith MR et al. 2004; Carr M and T Nielsen 2015). While some studies found a relative predominance of negative emotions, such as fear and anxiety, in dreams (Merritt J et al. 1994; Roussy F et al. 2000), others reported a balance of positive and negative emotions (Schredl M and E Doll 1998), or found that joy and emotions related to approach behaviors may prevail (Fosse R et al. 2001; Malcolm-Smith S et al. 2012). When performing a lexicostatistical analysis of large datasets of dream reports, a clear dissociation between dreams containing basic, mostly fear-related, emotions and those with other more social emotions (e.g. embarrassment, excitement, frustration) was found, highlighting distinct affective modes operating during dreaming, with fear in dreams representing a prevalent and biologically-relevant emotional category (Revonsuo A 2000; Schwartz S 2004). Thus, if fear-containing dreams serve an emotion regulation function, as hypothesized by the theoretical models, the stronger the recruitment of fear-responsive brain regions (e.g. amygdala, cingulate cortex, insula; see Phan KL et al. 2002) during dreaming, the weaker the reaction of these same regions to actual fear-eliciting stimuli during wakefulness should be. This compensatory or homeostatic mechanism may also be accompanied by an enhanced recruitment of emotion regulation brain regions (such as the medial prefrontal cortex, mPFC, which is implicated in fear extinction) during wakefulness (Quirk GJ et al. 2003; Phelps EA et al. 2004; Yoo SS *et al.* 2007; Dunsmoor JE et al. 2019).

Here, we collected dream reports and functional brain measures using high-density EEG (hdEEG) and functional MRI (fMRI) across two studies to address the following questions: (i) do emotions in dreams (here fear-related emotions) engage the same neural circuits as during wakefulness, and (ii) is there a link between emotions experienced in dreams and brain responses to emotional stimuli during wakefulness. By addressing these fundamental and complementary topics, we aim at clarifying the grounding conditions for the study of dreaming as pertaining to day/night affective homeostasis.

## METHODS

### Study 1: Neural correlates of fear in dreams

#### Participants

Eighteen healthy participants were included in Study 1 (4 males, age 39.77 ± 13.12 years, 25-63 [mean ± SD, range]). From these 18 participants, twelve (N=12) were used for the analysis of fear vs. no fear conditions in NREM sleep (N2 stage), while eight (N=8) were used for the analysis of fear vs. no fear conditions in REM sleep. All participants had no history of neurological or psychiatric disorder. Signed informed consent was obtained from all participants before the experiment, and ethical approval for the study was obtained from the University of Wisconsin-Madison Institutional Review Board.

#### Procedure

##### Serial awakenings during sleep

Dream sampling during sleep was accomplished using the ‘serial awakening’ method, as described in detail elsewhere (Siclari F *et al.* 2017) (Fig. 1A). In brief, participants were awakened several times during the night while sleeping and were asked to describe *‘the last thing going through your mind prior to the alarm sound’*, and then underwent a structured interview via intercom. Among other questions, they were asked to name any specific emotion that they experienced, and to report the presence/absence of fear or anxiety. Awakenings were performed at intervals of at least 20 minutes in N2 sleep or REM sleep using an alarm sound. Participants must have been asleep for a minimum of 10 minutes and must have been in a stable sleep stage for a minimum of 5 minutes before any experimental awakening was triggered.

**Figure 1.**
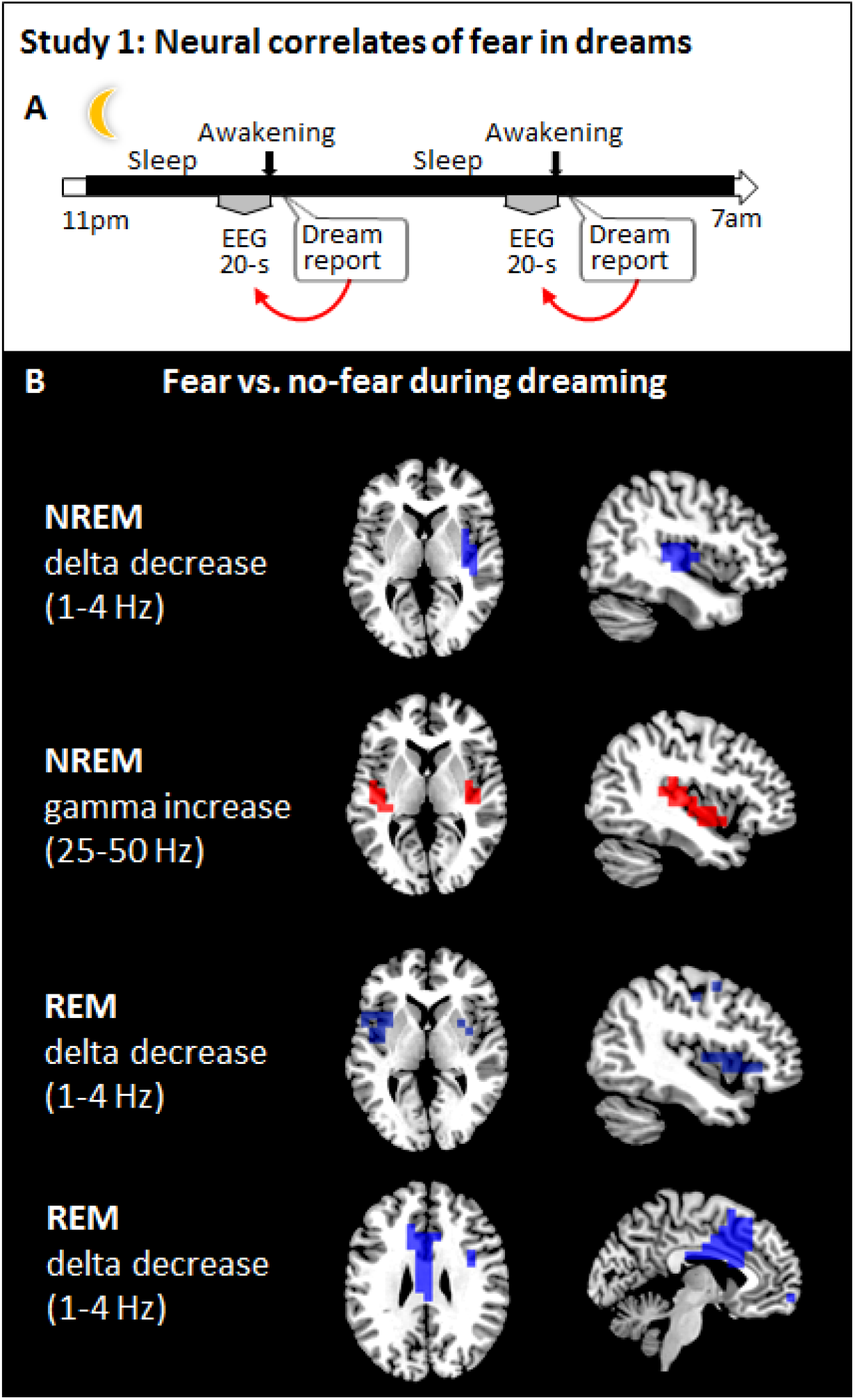
**A**. Participants under hdEEG were awakened several times while sleeping in the sleep laboratory and were asked to report “the last thing going through their [your] mind prior to the alarm sound” and then underwent a structured interview via intercom. Participants were also asked whether they felt fear. Twenty-second epochs of EEG recording prior to each awakening were then sorted as a function of the presence or absence of fear in the dream. **B.** Brain maps showing modulations of delta and gamma power during N2 sleep and delta power during REM sleep when comparing trials with fear to those without fear. Only significant differences at the p<.05 level, obtained after correction for multiple comparisons, are shown at the source level (two-tailed, paired t-tests, TFCE corrected).

#### EEG recordings

Recordings were made at the University of Wisconsin (Wisconsin Institute for Sleep and Consciousness) using a 256-channel high-density EEG (hdEEG) system (Electrical Geodesics, Inc., Eugene, Ore.) combined with Alice Sleepware (Philips Respironics, Murrysville, PA). Additional polysomnography channels were used to record and monitor eye movements and submental electromyography during sleep. Sleep scoring was performed over 30s epochs according to standard criteria (Iber C et al. 2007).

#### EEG Preprocessing

The EEG signal was sampled at 500 Hz and band-pass filtered offline between 1 and 50 Hz. The EEG data were high-pass filtered at 1Hz instead of lower frequencies as there were sweating artifacts in some of the participants which caused intermittent high-amplitude (>300uV) slow frequency oscillatory activity around 0.3 Hz. Noisy channels and epochs containing artifactual activity were visually identified and removed. To remove ocular, muscular, and electrocardiograph artifacts, we performed Independent Component Analysis (ICA) using EEGLAB routines (Schenck CH et al. 1993; Jung TP et al. 2000). The previously removed noisy channels were interpolated using spherical splines (EEGLAB). Finally, EEG data was referenced to the average of all electrodes.

#### EEG Signal analysis

##### Source localization

The cleaned, filtered and average-referenced EEG signal corresponding to the 20s before the alarm sound was extracted and analysed at the source level. Source modelling was performed using the GeoSource software (Electrical Geodesics, Inc., Eugene, Ore.). A 4-shell head model based on the Montreal Neurological Institute (MNI) atlas and a standard coregistered set of electrode positions were used to construct the forward model. The source space was restricted to 2447 dipoles in 3-dimensions that were distributed over 7×7×7 mm cortical voxels. The inverse matrix was computed using the standardized low-resolution brain electromagnetic tomography (sLORETA) constraint. A Tikhonov regularization procedure (λ=10^−1^) was applied to account for the variability in the signal-to-noise ratio (Pascual-Marqui RD 2002). We computed spectral power density using the Welch’s modified periodogram method (implemented with the *pwelch* function in MATLAB (The Math Works Inc, Natick, MA) in 2s Hamming windows (8 segments, 50% overlap) to decompose the source signals into frequency bands of interest before taking the norm across dimension to produce a single power value for each dipole.

##### Statistical Analysis

Statistical analysis was carried out in MATLAB. To compare brain activity between trials with fear and those without, source-space power was averaged within standard frequency bands (Delta: 1-4Hz, Theta: 5-8Hz, Alpha: 8.5-12Hz, Sigma: 12.5-17Hz, Beta: 17.5-24Hz, Gamma: 25-50Hz). We then averaged the power values within trials with fear and those trials without fear for each participant and for each frequency band separately. Group level analyses used paired two-sample t-tests (two-tailed) between the fear and no fear conditions, performed separately for each frequency band, and thresholded at corrected p< 0.05 using non-parametric threshold-free cluster enhancement (TFCE) (weighing parameters E=0.5 and H=2) (Mensen A and R Khatami 2013).

### Study 2: Modulation of brain responses to aversive stimuli during wake as a function of fear in dreams

#### Participants

A total of 127 healthy individuals (45 males, age 22.00 ± 3.15 years, 18-37 [mean ± SD, range]) participated in Study 2. All participants had no history of neurological or psychiatric disorder. Signed informed consent was obtained from all participants before the experiment, and ethical approval for the study was obtained from the University of Geneva (Switzerland) and University of Liège (Belgium). Among these 127 participants, 13 participants did not report any dream in their dream diary before the experimental session (see below). For the fMRI analyses, we also excluded 25 participants who were presented with emotional (and neutral) words, unlike all other participants who saw emotional pictures. The final group of 89 participants included 58 females, 21.5 ± 2.4 (mean ± SD) year-old. Participants took part in one of 3 different experiments (Exp. 1: N=28, age 21.36 ± 2.70 years, 16 men, University of Geneva; Exp. 2: N=19, age 22.16 ± 2.48 years, 19 men, University of Geneva, Exp. 3: N=42, age 21.31 ± 2.08, 23 men, University of Liège). All participants filled out the same questionnaires on sleep quality (PSQI, Tzourio-Mazoyer N et al. 2002), daytime sleepiness (Epworth sleepiness scale, ESS, Pascual-Marqui RD 2002), depression (Beck Depression Scale, BDI, Mensen A and R Khatami 2013), anxiety (State-Trait Anxiety Inventory Trait, STAI-T, Spielberger CD et al. 1970), and also kept the same sleep and dream diary. These results correspond to a secondary analysis because part of the data was already presented elsewhere (Sterpenich V, C Piguet, et al. 2014; Sterpenich V et al. 2017), yet without exploring any of the dream measurements. Note that the sleep and dream diaries were designed specifically for this analysis and the stimuli of the three different studies correspond to similar visual items (emotional and neutral pictures, see below), eliciting similar changes in emotional arousal and in local brain activition (Fig. S1).

#### Collection and analysis of dream data

During the week preceding the fMRI session, participants were asked to fill out a sleep and dream diary at home (Fig. 2A). Every morning, they responded to a mini-questionnaire about the content of the dreams they may have had during the preceding night, and had the possibility to write down their dreams in more details. Among other questions in the mini-questionnaire, participants were asked to report the presence/absence of specific emotions in their dreams (anger, disgust, confusion, embarrassment, fear, sadness, joy, frustration). Note that joyful dreams may be slightly over-represented because joy was the only positive emotion among the emotions assessed. We computed the percentage of nights with dreams containing specific emotions (for example, 3 nights with dreams containing fear over a total of 5 nights with dreams leads to a percentage of fear in dreams of 60%). We entered the percentage for each emotion from each participants (8 values by participant) into a Principal Component Analysis (PCA) to i) identify main affective components (or dimensions) according to variance explained and ii) characterize each participant by component (or factor) scores, which we then used as regressors in an fMRI design matrix. Specifically, here we used the individual scores on the second PCA component that contrasted basic negative emotions, in particular fear, to non-basic social negative emotions (such as embarrassment and frustration) (see Schwartz S 2004 for a similar data structure). Because of the imbalanced number of options for negative (n=7) vs. positive (n=1) emotions in the original data, we did not investigate the first PCA component. Indeed, and unsurprisingly, the latter contrasted the only positive emotion to all other (negative) emotions.

**Figure 2.**
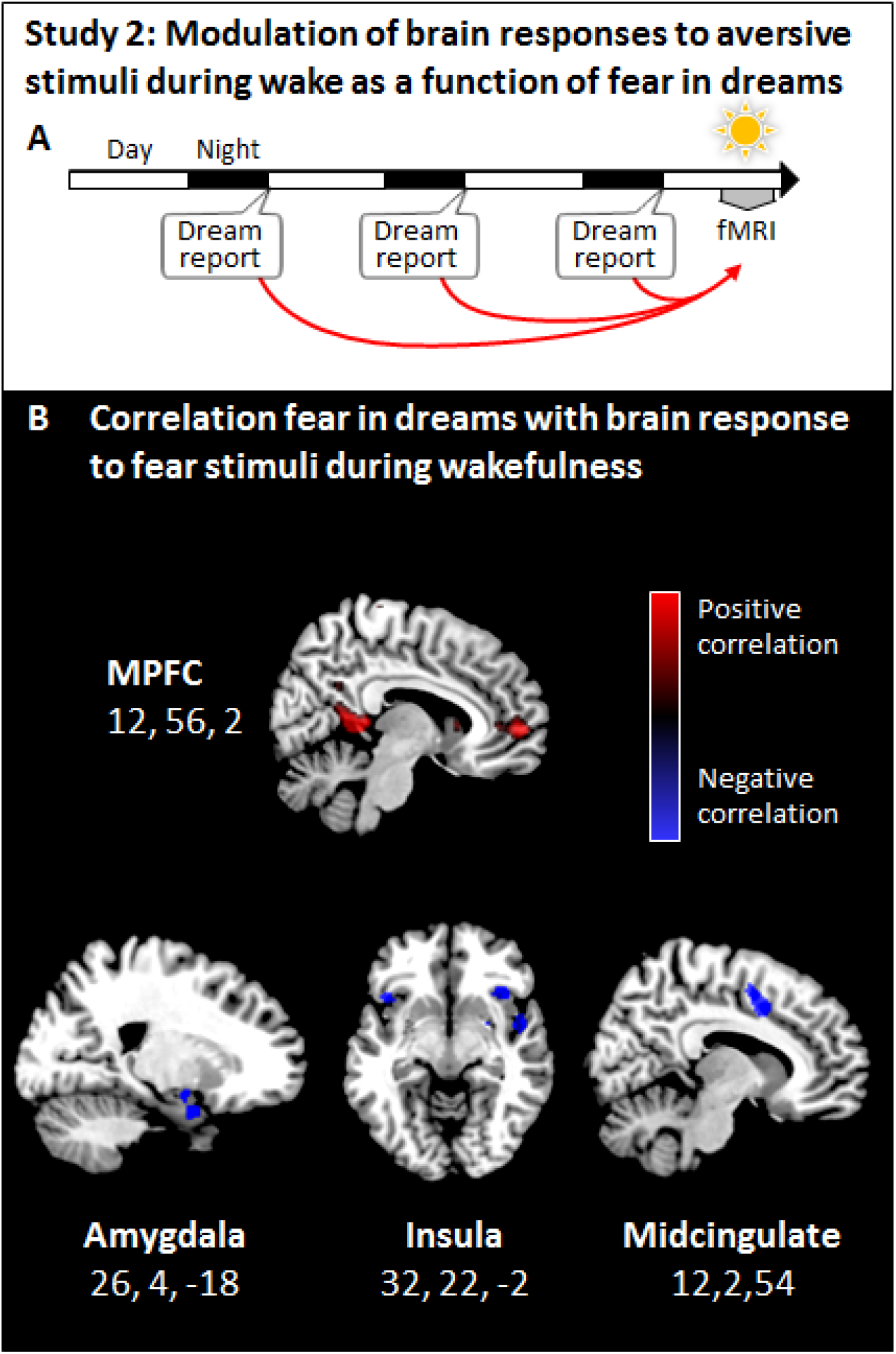
**A.** During one week, participants filled a dream diary at home, and their responses to aversive stimuli were assessed using fMRI. To test for a link between fear in dreams and brain responses to fear-eliciting stimuli during wakefulness, the individual propensity to experience fear in dreams (second PCA component, Table S1) was used as a regressor in a whole-brain analysis. **B.** Responses of the medial prefrontal cortex to aversive stimuli were greater in those individuals who frequently experienced fear in dreams (top panel), while activation of the amygdala, insula, and midcingulate cortex decreased in the same individuals (middle and bottom panels). Significant whole-brain regression results are displayed on the mean structural image.

#### Functional MRI session

##### Emotional tasks

Data from three different fMRI experiments were included in the analysis. Two of these three sets of data have already been reported elsewhere, but none of these former publications concerned emotions in dreams (Sterpenich V, C Piguet*, et al.* 2014; Sterpenich V *et al.* 2017). Common to these three experiments was that participants were exposed to aversive and neutral images, and that dream data were collected using the exact same instructions and dream diary. Below, we briefly describe the task used in each experiment, focusing only on those aspects that are relevant to the purpose of the present work.

In Experiment 1, participants were presented with conditioned (aversive) and unconditioned faces. Stimuli were presented in an intermixed, random order, one at a time (2.5 s each) followed by a varying interval (ISI: 4–5.5 s, mean=4.75) (see Sterpenich V, C Piguet*, et al.* 2014 for more details). In Experiment 2, 60 aversive, 60 funny and 60 neutral pictures were presented to participants. Each picture was displayed for 3s, preceded by a fixation cross of 1 s. Participants had a maximum of 2s to judge the valence of each stimuli on a 7 point scale (from −3: very negative, 0: neutral, +3: very funny). In Experiment 3, participants saw 90 faces displaying a negative expression and 90 faces with a neutral expression (see Nielsen T 2017 for more details). Each face was displayed for 3 s, and then participants judged it for emotional valence and arousal. Each trial started with a fixation cross for 1.5-s duration and ended as soon as participants responded, resulting in a jitter between trials (range 5.03-12.1 s). For the analysis of each experiment, we compared activity elicited by the presentation of aversive vs. neutral stimuli.

All visual stimuli were presented on a back projection screen inside the scanner bore using an LCD projector, which the participant could comfortably see through a mirror mounted on the head coil. Responses were recorded via an MRI-compatible response button box (HH-1 × 4-CR, Current Designs Inc., USA).

##### Pupillary size

During all fMRI sessions, eye movements and pupil diameter were measured continuously using an MRI-compatible long-range infrared eye tracking system (Applied Science Laboratories, Bedford, MA, USA; sampling rate: 60 Hz). Pupil size variation was used as an index of emotional arousal during the tasks (Bradley MM *et al.* 2008). Pupillary responses were analyzed during epochs of 5 s following the onset of the presentation of pictures. For each epoch, baseline pupil size was estimated as the average pupil measurement during the second preceding the presentation of the picture, and was then subtracted from all values of this epoch. Trials with more than 30% of signal loss were discarded. The pupillary values were z-scored for each task to take into account the difference in luminance of the different pictures and background. For each trial type (aversive and neutral) and for each task, we analyzed the mean signal value over the 5 s epochs using a t-test (Fig. S1A). Data from 50 participants were discarded because of poor quality of the recording or technical problems, and finally data from 77 participants were included in this analysis. The mean pupil diameter value for aversive stimuli was subtracted from that for neutral stimuli for each participant to obtain an individual physiological emotional response to fear-eliciting stimuli. To test for a potential link with emotions in dreams, we correlated these individual pupil reactivity values with the frequency of fear in dream.

##### MRI acquisition

For Experiments 1 and 2, MRI data were acquired on a 3 T whole body MR scanner (Tim Trio, Siemens, Erlangen, Germany) using a 12-channel head coil. For Experiment 3, data were acquired on a 3T head-only magnetic resonance scanner (Allegra, Siemens, Erlangen, Germany). Functional images were acquired with a gradient-echo EPI sequence with the following parameters. For Experiment 1: repetition time (TR): 2200ms, echo time (TE): 30ms, flip angle (FA): 85°, field of view (FOV): 235 mm, 36 transverse slices, voxel size: 1.8 × 1.8 × 3.4 mm. For Experiment 2: TR: 2100 ms, TE: 30 ms, FA: 80°, FOV: 205 mm, 36 transverse slices, voxel size: 3.2 × 3.2 × 3.8 mm. For Experiment 3: TR: 2460 ms, TE: 40 ms, FA: 90°, FOV: 220 mm, 32 transverse slices, voxel size: 3.4 × 3.2 × 3.4 mm. T1-weighted structural images were also acquired in each participant.

##### MRI analysis

Functional volumes were analyzed by using Statistical Parametric Mapping 8 (SPM8; www.fil.ion.ucl.ac.uk/spm/software/spm8) implemented in Matlab (The MathWorks Inc, Natick, Massachusetts, USA). Functional MRI data were corrected for head motion, slice timing and were spatially normalized to an echo planar imaging template conforming to the Montreal Neurological Institute (MNI) template (voxel size, 3 × 3 × 3 mm). The data were then spatially smoothed with a Gaussian kernel of 8 mm full width at half maximum (FWHM). For each participant, we used a General Linear Model (GLM) approach to estimate brain responses at every voxel, and computed the main contrast of interest (aversive vs. neutral). For each experiment, any other conditions (e.g. positive items in Exp. 2) were modeled as variable of no interest. Movement parameters estimated during realignment were also added as regressors of no interest. The resulting individual maps of t-statistics (the contrast images for each individual) were then used in second-level random-effects analyses. We used one-sample t-tests for testing common effects of aversive vs. neutral stimuli. For the regression analysis, individual scores on the second PCA component (higher for more fearful basic negative emotion in dreams; see Collection and analysis of dream data section) were used as a regressor at the group level for the contrast aversive vs. neutral stimuli. Statistical inferences were corrected for multiple comparisons according to the Gaussian random field theory at p<0.05 Family wise error (FWE) corrected i) on the entire volume or ii) using correction within predefined anatomical regions (small volume correction, SVC), including the amygdala, insula, and the midcingulate cortex using the toolbox Anatomy (Tzourio-Mazoyer N *et al.* 2002).

## RESULTS

### Study 1: Identifying the neural correlates of fear in dreams

In Study 1, we performed serial awakenings in 18 participants recorded throughout the night with hdEEG (256 channels) and identified brain regions activated prior to awakenings from a dream containing fear (vs. without fear; Fig. 1A). We only analyzed the EEG data from those participants who reported at least one dream containing fear and one without fear, within the same recording night and for a given sleep stage (N2 or REM). We conducted spectral analyses for the 20-seconds EEG epochs preceding each awakening and compared epochs associated with the presence of fear in dreams with those without fear. We found significant modulations in the delta (1-4 Hz) and gamma (25-50Hz) ranges, which we interpreted in terms of underlying local neuronal population activity as follows. Increased (respectively decreased) low frequency power in the delta range (<4Hz) corresponds to neuronal inhibition (respectively activation) (Tononi G and M Massimini 2008; De Gennaro L *et al.* 2011; Pigorini A et al. 2015), while increased high EEG frequencies in the gamma range reflect increases in neuronal firing (Steriade M et al. 1996; Le Van Quyen M et al. 2010) and positively correlate with local BOLD fluctuations (Murta T et al. 2015).

For NREM, we obtained a total of 79 awakenings in N2 sleep from 12 participants (average per night 6.58 ± 2.13 [mean ± SD, range]). Of these awakenings, 57 yielded reports of dream experience (from which 18 without recall of any content), while 22 yielded no report. Fear was present in 14 reports (average per subject 1.16 ± 0.37 [mean ± SD, range]) and absent in 25 reports (average per subject 2.08 ± 1.03 [mean ± SD, range]). N2 reports with presence of fear (compared to without fear) were associated with decreased delta power (1-4Hz) in the right insula, and increased gamma power (25-50Hz) in the bilateral insular cortex (Fig. 1B). No significant changes were found for other frequency bands.

For REM sleep, we obtained a total of 32 awakenings from 8 participants (average per night 4.00 ± 0.86 [mean ± SD, range]). Of these awakenings, 28 yielded reports of dream experience (from which 1 without recall of any content), while 4 yielded no report. Fear was present in 12 reports (average per subject 1.50 ± 0.50 [mean ± SD, range]) and absent in 15 reports (average per subject 1.87 ± 1.05 [mean ± SD, range]). REM reports with presence of fear compared to those without fear were associated with decreased delta power (1-4Hz) in the bilateral insula and midcingulate cortex, thus partly replicating the results from N2 (Fig. 1B). No significant changes were found for other frequency bands. Although EEG source reconstruction should be considered with caution, the present data also suggest that experiencing fear in NREM and REM dreams could involve different portions of the insula, with slightly more anterior dorsal insula activity during REM (Fig 1B). Together these results demonstrate that the occurrence of frightening dreams coincided with increased activation of insula cortex during both NREM and REM sleep, and of the midcingulate cortex during REM sleep.

### Study 2: Linking awake brain responses to aversive stimuli and fear in dreams

After we established that fear in dreams implicated the insula and midcingulate cortex known to contribute to the processing of aversive stimuli during wakefulness (Study 1), we asked whether frequently experiencing fear in dreams related to individual autonomic and neural sensitivity to fear during wakefulness. To this end, we collected a large amount of dream reports from 127 participants and correlated individual scores on a fear component (see below) with pupillary responses and fMRI responses to aversive stimuli during wakefulness recorded in the same participants (Fig. 2A). All participants filled the same sleep and dream diary over 7.38 (± 4.90; mean ± SD) nights with the same instructions before an MRI session (with a minimum of 3 nights). From the sleep diary, participants reported a mean sleep duration of 7.93 (± 0.76 SD) hours per night and a sleep quality of 7.02 (± 0.97 SD; on a 10-point scale from 0 - very bad sleep to 10 - very good sleep quality). From the dream diary, an average of 5.47 nights (± 3.63 SD) were associated with the feeling of having dreamt, and participants answered specific questions related to the content of their dreams after 3.75 (± 2.66 SD) nights. In particular, they were asked to indicate whether an emotion was present/absent for 8 distinct emotions (see Table S1). We performed a principal component analysis (PCA) on these data and found that the variance in emotion ratings was best explained by two main components. The first component (20.39% of the variance) distinguished between negative and positive (here “joy”) emotions, thus representing emotional valence. The second component (16.40% of the variance) contrasted emotions related to basic negative emotions (fear, anger, sadness, disgust), with fear having the strongest contribution, and those related to non-basic, social negative emotions (i.e., embarrassment, confusion, frustration; see (Schwartz S 2004) for a similar finding). The predominance of fear reported in dreams provides a first confirmation of our initial hypothesis about the expression of this emotion during sleep. Accordingly, we shall call this second “fear component” in the remainder of the manuscript. We did not use the first component in our fMRI analysis primarily because it does not directly address our main hypothesis about fear, and because of the imbalance between the number of negative and positive emotions participants had to choose from (7 vs. 1; Table S1), as also captured by the PCA. The individual PCA scores (or loadings) for the fear component did not correlate with sleep quality (PSQI, R^2^ <0.001, P=0.51), sleep duration (sleep diary, R^2^=0.005, P=0.44), sleepiness (ESS, R^2^=0.02, P=0.12), depression (BDI, R^2^=0.008, P=0.34), anxiety (STAI-T, R^2^=0.001, P=0.77), or frequency of dreaming (percentage of nights with dreams, R^2^=0.003, P=0.54). This pattern of results supports that the PCA fear component might represent an individual affective measure that is largely independent of other sleep-related variables and waking affective dimensions.

Study 2 aimed at establishing whether neurophysiological responses to fear-eliciting emotions during wakefulness correlated with fear in dreams. We first analyzed changes in pupillary size recorded during the presentation of aversive (vs. neutral; see Methods; Fig. S1A) stimuli as an index of emotional arousal response (Bradley MM et al. 2008), and showed that this index correlated with the individual PCA scores for the fear component (Spearman’s rho=0.25, P=0.034). We next tested whether we could observe a similar relationship between emotions in dreams and brain responses collected during the presentation of the same aversive stimuli (vs. neutral). Using fMRI data collected on 89 participants (from 3 experiments, see Methods for more details), we first confirmed that fear-eliciting stimuli activated a set of expected brain regions, including the amygdala, the insula, and occipital regions (Table S2, Fig. S1B). Then, we used the individual PCA scores for the fear component as a regressor in a whole-brain regression analysis (for the contrast aversive vs. neutral stimuli, see Methods). This analysis revealed that participants with a high propensity to experience fear in dreams had increased activation of the medial prefrontal cortex cortex (Fig. 2B, Table S3) when facing aversive stimuli while awake. Conversely, and as predicted by theoretical models (see Introduction), the same analysis yielded negative correlations with activity in the right amygdala, right insula, and midcingulate cortex (Fig. 2B). Taken together, these results show that individuals who reported a high prevalence of fear-related emotions in their dreams had stronger fear inhibition during wakefulness. Critically, associated neural decreases implicated the insula and midcingulate, which were both strongly activated during fearful dreams in Study 1, thus further supporting reciprocal links between sleep and wake emotional functioning.

## DISCUSSION

Here we investigated the neural correlates of fear in dreams and their relation to brain responses to threatening stimuli during wakefulness. In Study 1, we found that experiencing fear (vs. no fear) in dreams was associated with the activation of the insula and midcingulate cortex (the latter during REM dreams; Fig. 1B), which were both also activated when experiencing fear during wakefulness (Study 2; Fig. S1B, Table S2), as also classically reported in previous research (Pereira MG et al. 2010; Alves FH et al. 2013; Casanova JP et al. 2016). We recently reported that specific dream contents—such as faces, places, movement, speech, and thoughts— engage similar cortical networks as during wakefulness (Perogamvros L et al. 2017; Siclari F et al. 2017). Here we show, for the first time to our knowledge, that a specific emotional state, fear, activated the insula and midcingulate during both dreaming and awake consciousness. The consistency of the present results across brain states and their correspondence with classical work on brain structures involved in fear are encouraging. Importantly, hdEEG (especially with 256 electrodes as in our study) can have sufficient accuracy in source localization (e.g. associating dreams containing faces with activation of the fusiform face area (Siclari F *et al.* 2017), even for the detection of signal originating from deep cerebral structures (Seeber M et al. 2019). However, combined EEG/fMRI studies are certainly needed to further elucidate the exact contribution of subcortical structures, such as the amygdala, especially during REM sleep (Maquet P *et al.* 1996; De Gennaro L *et al.* 2011; Corsi-Cabrera M *et al.* 2016).

The insula, especially its anterior part, may contribute to social–emotional experience and associated visceral states, possibly giving rise to conscious feelings (Critchley HD et al. 2004; Chang LJ et al. 2013), and participates in the emotional response to distressing cognitive or interoceptive signals (Reiman EM et al. 1997). Insula activation during dreaming could thus reflect the integration of internally-generated sensory, affective and bodily information culminating in a subjective feeling of danger, as we further discuss below. During REM sleep, we also found an activation of the midcingulate cortex, a region known to be critically involved in behavioral/motor responses to dangers (Pereira MG *et al.* 2010). Because REM sleep is characterized by activation across sensory and motor cortices (Schwartz S and P Maquet 2002), while muscle atonia prevents the overt expression of motor behaviors, this sleep stage could provide a well-suited physiological condition for the (re)activation of threatening situations with associated emotional and motor reactions.

Based on the results from Study 1, which suggested an anatomo-functional correspondence between fear in dreams and wakefulness, we then asked whether frequently experiencing fear in dreams may relate to the individual’s neurophysiological sensitivity to fear during wakefulness. In Study 2, we analyzed awake fMRI response to aversive stimuli as a function of whether participants reported a high incidence of fear in their dreams. Here we thus considered fear in dreams as an individual trait, which we determined based on the analysis of a large dataset of dreams. Of note, this measure of fear in dreams did not correlate with depression, anxiety, sleep quality, and the frequency of dream recall (all p>0.05), which supports the specificity of fear in dreams with respect to other classical dimensions of mood or sleep/dreams. We found decreased activity in the insula, amygdala and midcingulate cortex, while activity of the mPFC cortex was increased (Fig. 2B). Along with the insula, both the amygdala and the cingulate cortex have been associated with fear and the perception of negative emotions during wakefulness (Phan KL *et al.* 2002), as we also demonstrated (Fig. S1B, Table S2). On the other hand, the mPFC is believed to regulate the response to threatening stimuli by modulating the activity of the amygdala (Quirk GJ *et al.* 2003; Phelps EA *et al.* 2004). Specifically, the mPFC exerts an inhibitory control on fear expression by decreasing amygdala output and has been associated with extinction learning (i.e., when a neutral conditioned stimulus that previously predicted an aversive unconditioned stimulus no longer does so, and conditioned response subsequently decreases) (Kalisch R et al. 2006; Herry C et al. 2010). Consistent with the proposal that dreaming may serve an emotion regulation function (Kramer M 1991; Hartmann E 1996; Cartwright R *et al.* 2006; Perogamvros L et al. 2013), participants who frequently (but not excessively, see below) experienced frightening dreams showed a stronger inhibition of the amygdala, potentially mediated by the mPFC. This interpretation is further supported by the pupillary results showing that participants who frequently reported fear in their dreams had reduced autonomic responses to aversive stimuli during wakefulness, suggesting a better ability to regulate defensive and alerting reactions to threatening signals in those individuals.

In the domain of sleep research (irrespective of dreaming), REM sleep was found to play a role in emotional memory consolidation (Nishida M et al. 2009; Goldstein AN and MP Walker 2014; Sterpenich V, C Schmidt, et al. 2014), especially fear memory consolidation (Pace-Schott EF et al. 2015) and successful fear/safety recall (Menz MM et al. 2016), while both NREM (Hauner KK et al. 2013; He J et al. 2015) and REM sleep stages (Pace-Schott EF *et al.* 2015; Menz MM *et al.* 2016) have been found to promote the retention and generalization of extinction learning. In addition, it was proposed that specifically REM sleep contributes to the attenuation of the emotional tone of waking-life memories (Walker MP and E van der Helm 2009; Vallat R et al. 2017). Importantly, total sleep deprivation may cause a reduction of mPFC control over the limbic system, resulting in an accentuation of emotional responses to negative stimuli (Yoo SS *et al.* 2007) and an impairment of extinction recall (Straus LD et al. 2017). We have previously suggested that these findings from sleep studies may putatively extend to or even depend on concomitant dreaming (Perogamvros L and S Schwartz 2012; Perogamvros L *et al.* 2013). In particular, the exposure to feared stimuli (objects, situations, thoughts, memories, and physical sensations) in a totally safe context during dreaming would thus resemble desensitization therapy (Levin R and TA Nielsen 2007). Besides, several studies have demonstrated that dreaming of negative waking-life experiences (e.g. divorce) contributes in a resolution of previous emotional conflicts and a reduction of next-day negative mood (Cartwright RD et al. 1984; Cartwright R *et al.* 2006). While this was not the objective of the present work, we would like to emphasize that the present data do not allow us to make any inference about whether one occurrence of one specific emotion in a dream influences emotional state or responsiveness on the following day.

Contrasting with this beneficial role of negative but benign dreams, recurrent nightmares, such as those observed in PTSD patients, might represent a failure of the fear extinction function of dreaming (Nielsen T and R Levin 2007; Nielsen T 2017). Thus, nightmare patients may be more prone to emotional dysregulation, as suggested by one recent study reporting decreased mPFC activity during the viewing of negative pictures in these patients (Marquis L et al. 2016). Furthermore, exerting ineffective emotional regulation strategies (e.g. fear suppression) and elevated anxiety during wakefulness may lead to increased excitability of negatively-loaded memories at sleep-onset or even during sleep (Schmidt RE and GH Gendolla 2008; Malinowski J 2017; Sikka P et al. 2018; Sikka P et al. 2019), namely in conditions where monitoring from the prefrontal cortex is reduced (Maquet P *et al.* 1996; Braun AR *et al.* 1997). Such disruption in the regulation of emotions during wakefulness and sleep has been proposed as a major contributing factor to insomnia (Wassing R et al. 2016).

Here, we experimentally show that, beyond sleeping, experiencing negative emotions *specifically* during dreaming is associated with better-adapted emotional responses during waking life. Study 2 combined data from three different experiments testing for brain responses to aversive stimuli (vs. neutral). This allowed us to include a very large set of participants, which is needed to exploit interindividual differences, as we do here. In all three experiments, dream reports were collected using the exact same instructions and same questionnaires. While we confirmed consistent fMRI and pupillary response results across the three experiments for the effects of aversive vs. neutral emotions, Study 2 yielded significant results in brain regions, for which we had strong theory-driven a priori. We therefore suggest that what may be perceived as a potential limitation (i.e. combining data from 3 experiments) may actually offer a better generalizability of the present findings to diverse waking threatening situations.

Taken together, across two complementary studies, we show opposing neural effects of fear experience in dreams and during wakefulness. These results support recent theoretical claims that dreaming (beyond sleep) benefits emotion regulation processes, by achieving a form of overnight affective simulation or recalibration (e.g. through extinction learning and generalization), which would foster adapted emotional responses to dangerous real-life events (Revonsuo A 2000; Nielsen T and R Levin 2007; Perogamvros L and S Schwartz 2012). Studying the role of positive emotions (e.g. positive social interactions) in dreams (especially in NREM dreams, (McNamara P et al. 2005) and their potential links with emotional brain responses during wakefulness may be needed to further corroborate or refine existing theoretical models. Finally, based on our results, we would like to suggest that future studies should address how sleep and dreaming may influence exposure and extinction-based therapies for affective disorders (Pace-Schott EF et al. 2018).

## ACKNOWLEDGMENTS

This study was supported by the Swiss National Science Foundation Grants 155120 (to L.P.) and 320030-159862 (to S.S.), NIH/NCCAM P01AT004952 (to G.T.), NIH/NIMH 5P20MH077967 (to G.T.), Tiny Blue Dot Inc. grant MSN196438/AAC1335 (to G.T.).

## AUTHOR CONTRIBUTIONS

VS, LP, GT, and SS designed the experiments, VS and LP conducted the experiments, VS, LP, and SS analyzed the data, VS, LP, GT and SS wrote the paper.

---

**Table S1:** PCA analysis of the emotions in dreams.

| Type of emotion | Percentage of emotion in dreams | PCA component 1<br>(correlation coefficient) | PCA component 2<br>(correlation coefficient) |
| --- | --- | --- | --- |
| Fear | 19.42 ± 27.38% | -0.40 | 0.36 |
| Anger | 15.63 ± 23.11% | -0.65 | 0.16 |
| Sadness | 9.65 ± 20.10% | -0.41 | 0.07 |
| Disgust | 5.15 ± 13.82% | -0.62 | 0.04 |
| Frustration | 13.21 ± 21.16% | -0.62 | -0.33 |
| Confusion | 18.05 ± 25.44% | -0.35 | -0.53 |
| Embarrassment | 8.31 ± 18.59% | -0.08 | -0.59 |
| Joy | 34.84 ± 32.41% | 0.22 | -0.64 |

**Table S2:** Functional MRI responses to aversive (vs. neutral) stimuli across the three fMRI experiments. Results are corrected for multiple comparisons using family-wise error correction at p<0.05 (FWE-corr) i) on the entire volume or ii) on predefined anatomical regions of interest (marked as ^+^).

| Name | MNI coordinates | Z-score | P<sub>FWE-corr</sub> |
| --- | --- | --- | --- |
| ***Aversive > Neutral*** |  |  |  |
| Amygdala | 20, -4, -16 | 4.51 | <0.001<sup>+</sup> |
| Anterior insula | -36, 26, -6 | 6.24 | <0.001 |
|  | 32, 16, -18 | 5.20 | 0.003 |
| Inferior frontal gyrus | -32, 14, -26 | 5.87 | <0.001 |
|  | 44, 32, -4 | 5.64 | <0.001 |
| Motor cortex | -42, 0, 46 | 4.95 | 0.010 |
| Anterior middle temporal gyrus | 50, -2, -24 | 6.74 | <0.001 |
| Posterior middle temporal gyrus | -54, -58, 10 | 6.16 | <0.001 |
|  | 52, -56, 14 | 6.06 | <0.001 |
| Thalamus | 0, -16, -4 | 6.45 | <0.001 |
| Superior temporal sulcus | 50, -18, 10 | 5.75 | <0.001 |
| Calcarine/lingual gyrus | -10, -78, 8 | 4.87 | 0.015 |
|  | 20, -60, 4 | 5.19 | 0.003 |
| Cuneus | 14, -100, 6 | 4.98 | 0.009 |
|  | -28, -86, -6 | 4.90 | 0.013 |
| Midcingulate cortex | 10, 18, 42 | 4.43 | 0.002<sup>+</sup> |
| Superior frontal sulcus | 42, 12, 24 | 5.04 | 0.007 |
| ***Aversive < Neutral*** |  |  |  |
| Medial PFC | 10, 44, -8 | 4.77 | 0.009 |
| Motor cortex | -52, -16, 50 | 4.91 | 0.013 |
| Angular gyrus | 50, -62, 46 | 4.99 | 0.009 |

**Table S3:** Whole-brain correlation between fMRI activity elicited by aversive (vs. neutral) stimuli during wakefulness and basic (fear) emotions in dreams (as captured by the score on the second PCA component). Results are corrected for multiple comparisons using family-wise error correction at p<0.05 (FWE-corr) i) on the entire volume or ii) on predefined anatomical regions of interest (marked as ^+^).

| Name | MNI coordinates | Z-score | P<sub>FWE-corr</sub> |
| --- | --- | --- | --- |
| ***Positive correlations*** |  |  |  |
| Medial PFC | 12,56,2 | 5.59 | 0.004 |
| ***Negative correlations*** |  |  |  |
| Amygdala | 26, 4, -18 | 3.48 | 0.008<sup>†</sup> |
| Amygdala/striatum | 22,0,-12 | 3.48 | 0.006<sup>†</sup> |
| Anterior insula | -30, 26, 4 | 3.33 | 0.058<sup>†</sup> |
|  | 32, 22, -2 | 3.65 | 0.019<sup>†</sup> |
| Midcingulate cortex | 12, 2, 54 | 4.68 | 0.034 |

**Figure S1.**
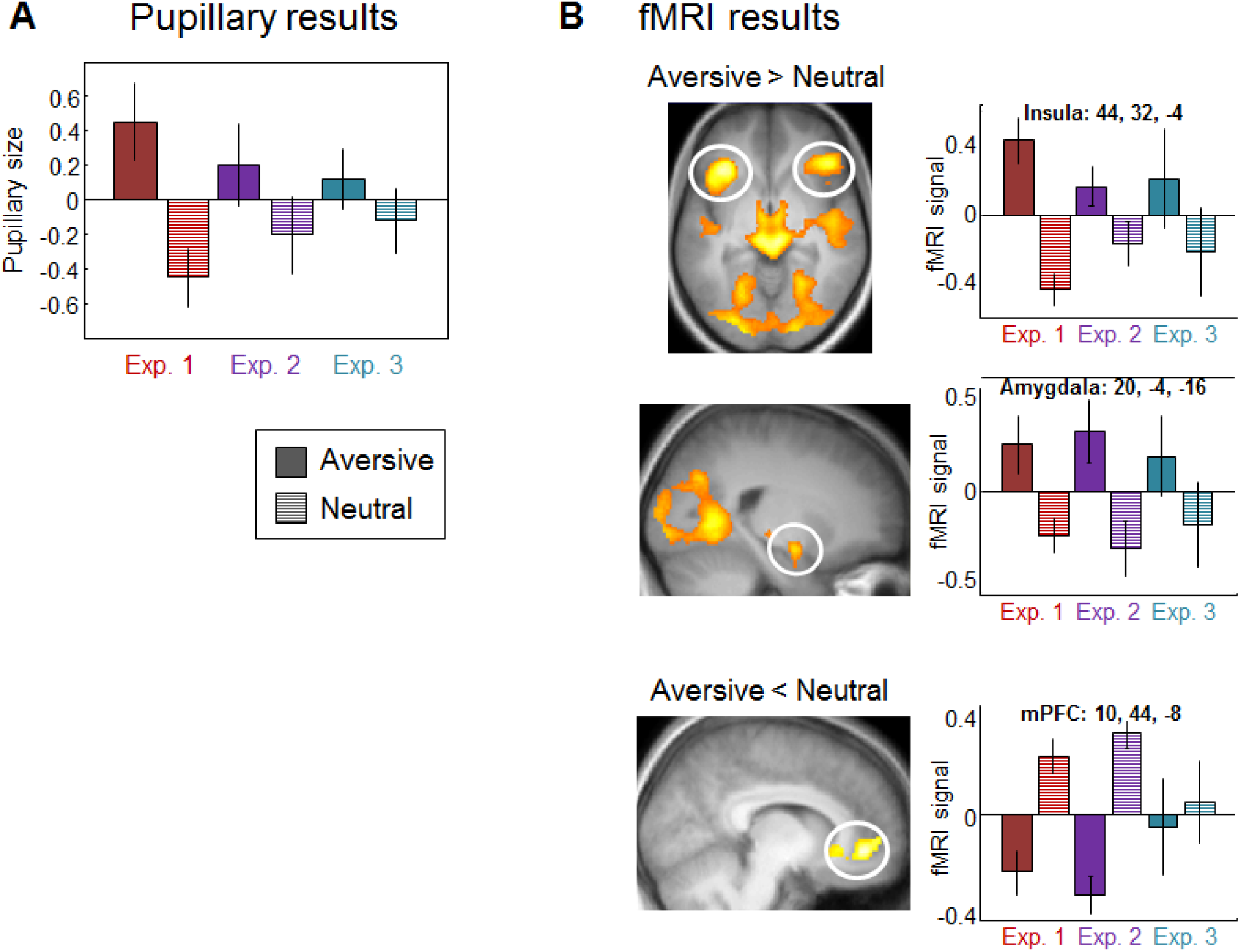
**A.** Pupil diameter change in response to aversive and neutral stimuli, showing a significant difference between aversive (plain color) and neutral (hatched) conditions for each experiment (Exp. 1: t=4.79, p<0.001; Exp. 2: t=5.15, p<0.001; Exp. 3: t=3.49, P=0.002). **B.** Regional changes in fMRI signal in response to aversive vs. neutral stimuli across the three fMRI experiments. Right panels show the parameters estimates for each experiment separately for aversive and neutral stimuli. For display purposes, whole-brain results are displayed on the mean structural image at P=0.001 uncorrected.

